# Identifying epigenetic biomarkers of established prognostic factors and survival in a clinical cohort of individuals with oropharyngeal cancer

**DOI:** 10.1101/679316

**Authors:** Ryan Langdon, Rebecca Richmond, Hannah R. Elliott, Tom Dudding, Nabila Kazmi, Chris Penfold, Kate Ingarfield, Karen Ho, Andrew Bretherick, Chris Haley, Yanni Zeng, Rosie M Walker, Michael Pawlita, Tim Waterboer, Sue Ring, Tom Gaunt, George Davey Smith, Matthew Suderman, Steve Thomas, Andy Ness, Caroline Relton

**Affiliations:** MRC Integrative Epidemiology Unit at the University of Bristol, UK; Population Health Sciences, Bristol Medical School, University of Bristol, UK; NIHR Bristol Biomedical Research Centre, University Hospitals Bristol and University of Bristol, UK; MRC Human Genetics Unit, Institute of Genetics and Molecular Medicine, University of Edinburgh, Western General Hospital, Crewe Road, EH4 2XU, Scotland, UK; Faculty of Forensic Medicine, Zhongshan School of Medicine, Sun Yat-Sen University, Guangzhou, China; Guangdong Province Translational Forensic Medicine Engineering Technology Research Center, Zhongshan School of Medicine, Sun Yat-Sen University, Guangzhou, China; Medical Genetics Section, Centre for Genomic and Experimental Medicine, Institute of Genetics and Molecular Medicine, University of Edinburgh, Edinburgh, EH4 2XU, UK; Centre for Cognitive Ageing and Cognitive Epidemiology, University of Edinburgh, Edinburgh, EH8 9JZ, UK; Infections and Cancer Epidemiology, German Cancer Research Center (DKFZ)

## Abstract

Smoking status, alcohol consumption and HPV infection (acquired through sexual activity) are the predominant risk factors for oropharyngeal cancer and are thought to alter the prognosis of the disease. Here, we conduct epigenome-wide association studies (EWAS) of these factors and ∼3-year survival using Illumina Methylation EPIC blood DNA methylation profiles from 409 individuals in the Head and Neck 5000 (HN5000) study. CpG site associations below our multiple-testing threshold (*P*_*Bonferroni*_ < 0.05) with both a prognostic factor and with survival were observed in four gene regions: *SPEG* (smoking), *GFI1* (smoking), *PPT2* (smoking), and *KHD3CL* (alcohol consumption). These were further analysed using 2-step Mendelian randomization to assess whether methylation may be a causal mediator of cancer survival. Evidence for mediation was observed only in the *SPEG* gene region, showing an association with decreased survival (mortality HR: 1.28, 95% CI: 1.14 to 1.43, *P:* 2.12×10^−05^). Replication in data from independent datasets, and from HN5000 participants with longer follow-up times is needed to confirm these findings.

## Introduction

Head and neck cancer (HNC) is the eighth most commonly diagnosed type of cancer, with over 12,000 new cases diagnosed in the UK in 2015 (1). Recently, oropharyngeal cancer (OPC), a subtype of HNC, has shown a significant increase in incidence in the UK. It has more than doubled between 1990 and 2006, with a further doubling since 2010 (2) and is affecting younger populations (<45 years old) with greater frequency (3). OPC shows poor survival rates, with the 5-year relative survival rate for the more recently diagnosed oropharyngeal cases (between 2009-2013) estimated to be around 55-60% (4).

Several lifestyle factors as well as viral infections have been implicated in altering both incidence and prognosis for OPC (5-7). Of particular importance for both incidence (5, 8, 9) and prognosis of OPC (10) are smoking, alcohol intake and HPV16 infection (via sexual contact, including that of oral sex). Smoking and, to a lesser extent, heavy drinking at the time of diagnosis are both associated with increased incidence and poor prognosis (10-12). Interestingly, HPV16 infection, while being a risk factor for OPC incidence, has been associated with better prognosis (13-15), with a population-based study conducted in Boston from December 1999 to December 2003 demonstrating that HPV16 infected cases showed improved overall survival compared to those without an infection (HR: 0.1; 95% CI: 0.02-0.4; N: 448) (16).

Epigenetic signatures can be measured using rapid high-throughput approaches and may serve as valuable prognostic markers for cancer (17). While several whole-genome methylation assays have been performed to define the DNA methylation signatures of tumour samples (18, 19), the ability to study cancers through non-invasive sampling of body fluids is a rapidly advancing development in cancer diagnostics and prognosis. In particular, biomarkers identified in blood hold promise as non-invasive prognostic tools and may potentially be used to direct treatment if shown to be informative proxies for cancer development and prognosis (20).

Ultimately, smoking, alcohol consumption, and HPV16 infection may influence blood DNA methylation patterns which, therefore, have the potential to act as novel exposure or prognostic indicators (21-23). Furthermore, as epigenetic changes are a hallmark process of cancer (24), DNA methylation patterns associated with cancer survival may provide insight into biologically relevant pathways. More specifically, these epigenetic changes may act as intermediates on the pathways by which exposures influence survival. For example, as viral infections are thought to play an important role in altering epigenetic processes (25-27), these may serve as a mechanism by which having a HPV16 infection might confer a protective effect on survival. However, distinguishing a causal mediating role of these epigenetic changes from other explanations, such as confounding and reverse causation, is challenging and requires techniques such as Mendelian randomization (MR) to strengthen causal inference (28-30). MR is an approach which uses genetic variants strongly associated with modifiable exposures to appraise the causal effect of an exposure on disease risk. This approach has been extended to interrogate the causal relationship with molecular intermediates such as DNA methylation (29, 30).

In the setting of a large prospective head and neck cancer cohort (the Head and Neck 5000 Study; HN5000), we profiled blood DNA methylation in 443 participants with oropharyngeal cancer close to the time of diagnosis and prior to treatment starting. We performed epigenome-wide association analyses (EWAS) of the main prognostic factors for oropharyngeal cancer (alcohol, smoking and HPV16 infection) as well as survival up to ∼3 years. We then assessed overlap between the DNA methylation profiles related to these prognostic factors and survival. Where there was evidence of a shared signal, we performed Mendelian randomization analysis to appraise the causal effect of DNA methylation in mediating the effect of these factors on survival.

## Results

Baseline characteristics of samples with epigenetic data, compared to all HNC and OPC samples in HN5000 are shown in **Table 1**. The proportion of those under the age of 60, and the proportion of those which are HPV16 E6 seropositive (an established biomarker of HPV-driven OPC) in OPC vs non-OPC HNC is notably greater. **Table 1** shows that those with OPC who were selected to have their methylation patterns typed were broadly representative of others with OPC in HN5000 with respect to exposure to prognostic factors, albeit not necessarily representative of HNC as a combined entity.

**Table 1.** Comparison of patient demographics in OPC samples selected for methylation data extraction, all samples in HN5000 identified as OPC, and all samples in HN5000

| Variable | OPC in HN5000 with methylation data and complete phenotype data (N=409) | OPC in HN5000 (N=1,909) | All HN5000 (all sub-types) (N=5,392) |
| --- | --- | --- | --- |
| ICD group (% oropharynx) | 100 | 100 | 35.4 |
| Sex (% female) | 27.0 | 21.9 | 27.2 |
| Age (% <60) | 58.4 | 52.4 | 42.7 |
| Smoking (% never smoked) | 27.1 | 28.0 | 24.6 |
| Alcohol (% non-drinker) | 25.9 | 26.6 | 28.4 |
| HPV16 E6 (% negative) | 33.3 | 32.3 | 72.0 |
| Survival (% died, prior to 30/09/2017) | 26.2 | 24.2 | 28.0 |

### Epigenome-wide association study of smoking

The single-site EWAS of ever vs never smokers revealed 52 CpG site associations annotated to 27 unique loci (P<5.7×10^−8^, Bonferroni-adjusted *P* < 0.05 for 862,491 tests) (**Figure 1**). CpG site cg05575921, which annotates to the *AHRR* gene region, was most strongly associated (P< 1.48×10^−40^) and showed the largest effect size of −29.5% difference in methylation between ever and never smokers. Forty-nine of the associated CpG sites had lower DNA methylation in ever smokers, with a mean difference in methylation of −8.3% (SD: 5.1%, range: −29.5% to −2.2%). The three remaining CpG sites had higher methylation in smokers, with a mean difference of 7.7% (SD: 4.2%, range: 4.7% to 12.6%). **Supplementary Table 1** provides the complete list of all CpGs that were differentially methylated below a multiple testing threshold of *P*:2.4×10^−7^ (the literature-reported alpha for the Illumina 450K BeadChip (31), a predecessor of the EPIC array, common in epidemiological literature, which can assay >450,000 CpG sites compared to >850,000 on the EPIC array). Of the results presented in this table, 37.5% (24/64 CpGs) were CpG sites present on the EPIC array but not its 450K predecessor.

**Figure 1.**
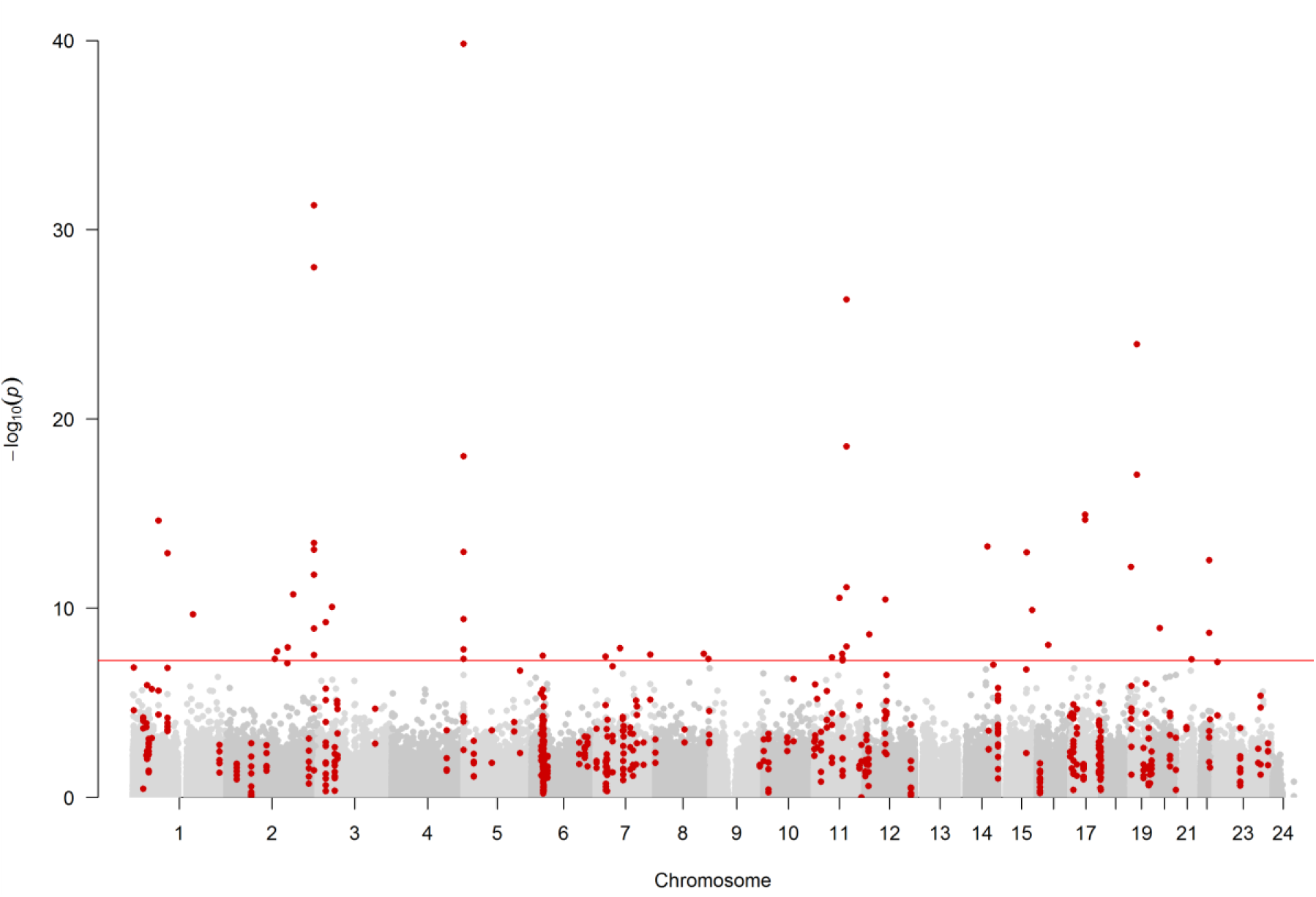
Manhattan plot of EWAS results from a comparison of ever vs. never smoking, showing CpG sites within DMRs in red. Each dot represents a single CpG site, plotting –log_10_(p) (y-axis) against the genomic position of the CpG site (x-axis). The horizontal red line is at P<5.7×10^−8^ and represents the value below which methylation was deemed to be significantly associated with smoking.

In the differentially methylated region (DMR) analysis of ever vs never smoking, 166 unique DMRs containing 617 measured CpGs and mapping to 156 gene regions were identified (**Figure 1**). The DMR with the strongest association contained 3 measured CpGs (cg21566642, cg01072057 and cg13903162) and was located at Chr2:233284661-233285290, an intergenic CpG island on 2q37.1 (P:1.13 ×10^−46^).

### Epigenome-wide association study of alcohol

The EWAS of alcohol consumption revealed 3 CpG site associations annotated to 3 unique genes (P<5.7×10^−8^) **(Figure 2**). The association with the smallest p-value was cg06690548 (P:8.3×10^−16^), annotating to the *SLC7A11* gene region. This CpG site also showed the largest effect size of −0.10% difference in methylation per unit of alcohol increase. All results below the 450K array multiple testing threshold of 2.4×10^−7^ are shown in **Supplementary Table 2.** Of the results presented in this table, 40% of the CpGs (2/5 CpGs) were present on the EPIC array but not it’s 450K predecessor.

**Figures 2.**
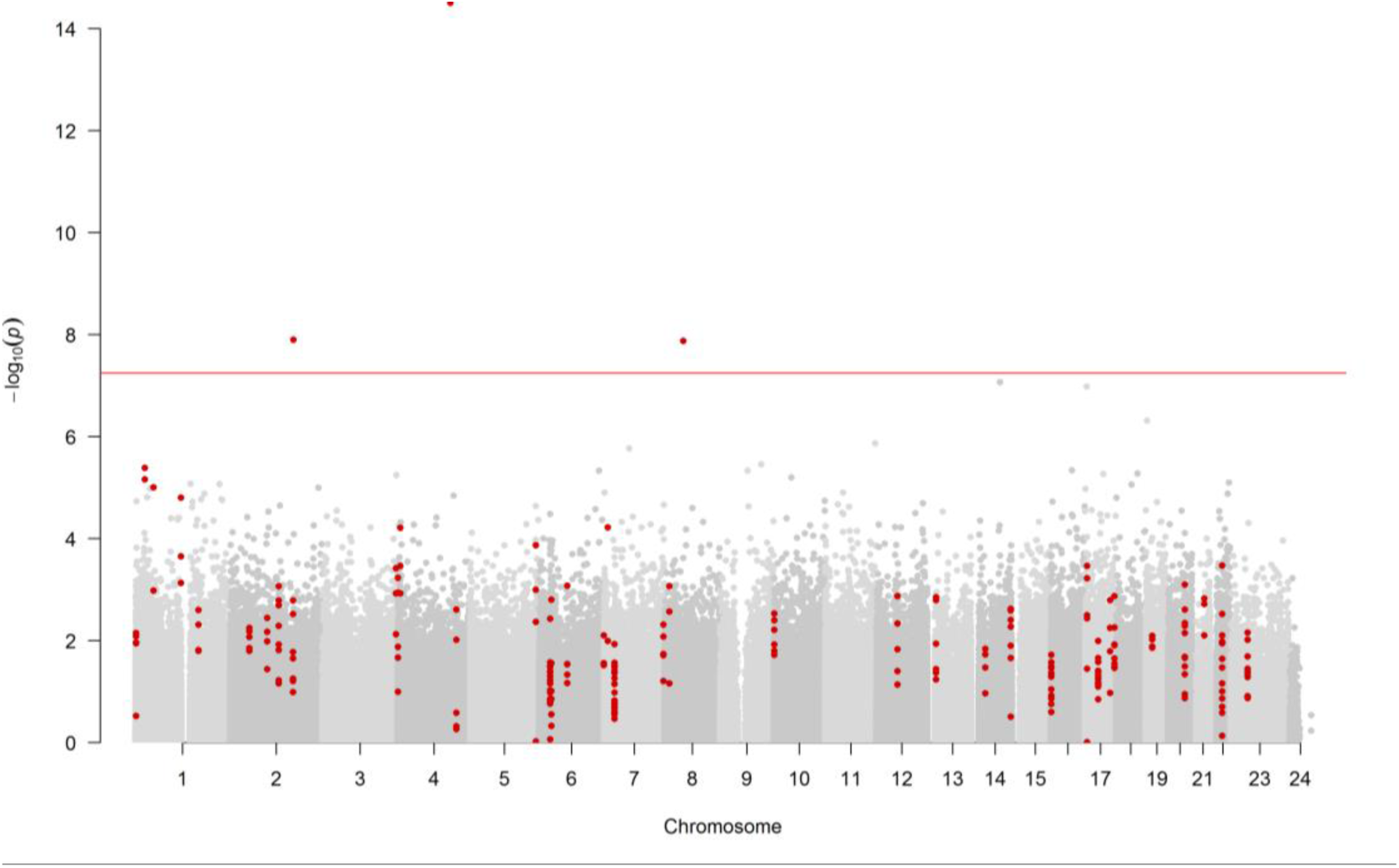
Manhattan plot of EWAS of alcohol consumption, showing CpG sites within DMRs in red. Each dot represents the EWAS result for a single CpG site, plotting –log_10_(p) (y-axis) against the genomic position of the CpG site (x-axis). The horizontal red line is at P<5.7×10^−8^ and represents the value below which CpG sites were considered to have good evidence of association with alcohol consumption.

In the DMR analysis of alcohol consumption, 40 unique DMRs containing 238 measured CpGs and mapping to 34 gene regions were identified (**Figure 2**). The DMR with the smallest P value was a region containing 2 CpGs (cg06690548 and cg13903162) found at Chr4:139162808-139163020 (P:1.45 ×10^−10^), annotating to the *SLC7A11* gene region.

### Epigenome-wide association study of HPV seropositivity

In the EWAS analysis of HPV16 E6 seropositivity, no CpGs passed our multiple testing p-value threshold (P<5.7×10^−8^) (**Figure 3**). At a suggestive threshold of 2.4×10^−7^, only 1 CpG site (cg26738437; *P:*1.3×10^−7^) was found, annotating to the *CCL16* gene. This probe is not found on the 450K array. Methylation at this site was on average 2.3% lower in HPV16 E6 seropositive participants when compared to controls.

**Figure 3.**
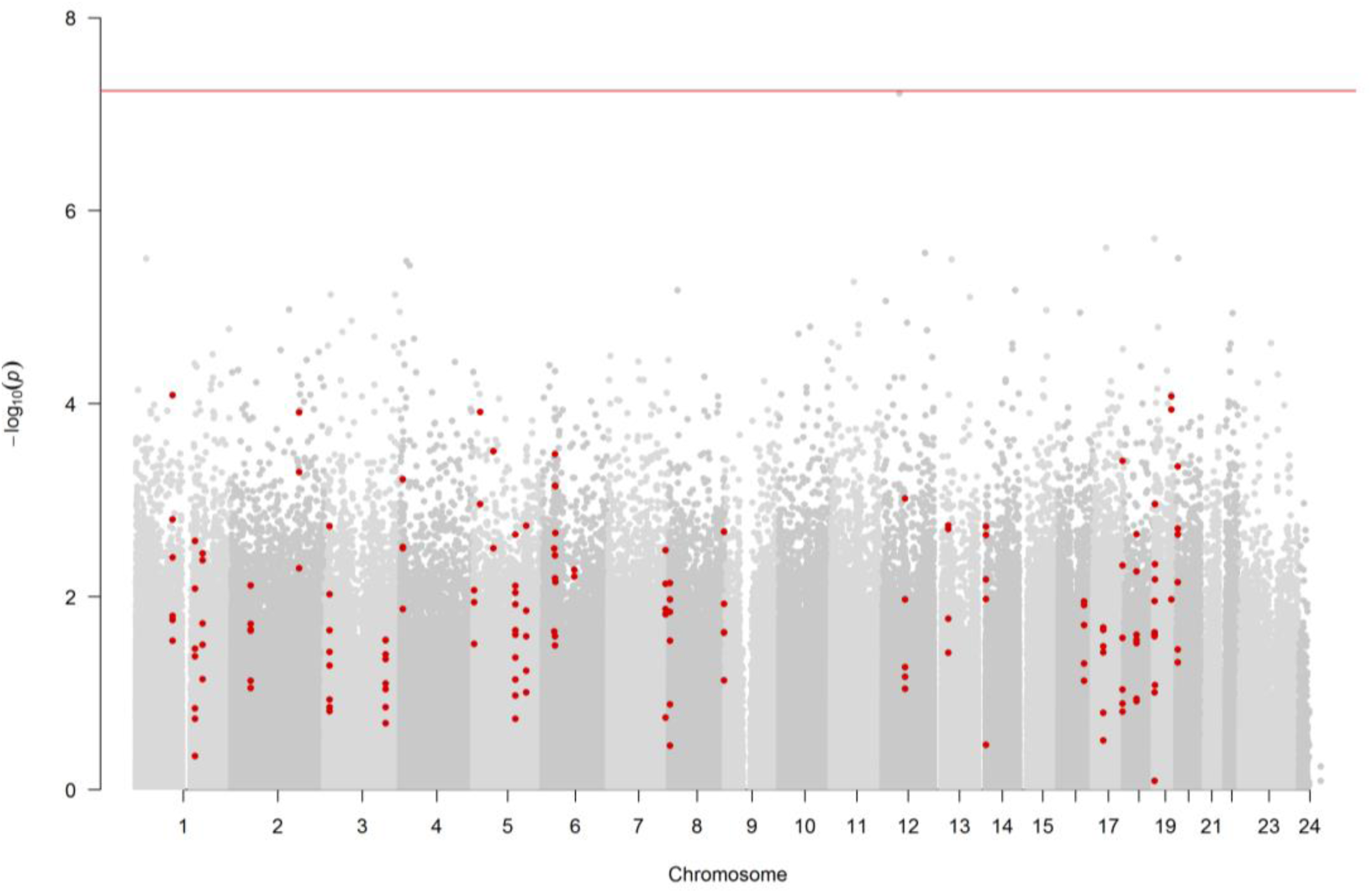
Manhattan plot of EWAS of HPV16E6 seropositivity, showing CpG sites within DMRs in red. Each dot represents the EWAS result for a single CpG site, plotting –log_10_(p) (y-axis) against the genomic position of the CpG site (x-axis). The horizontal red line is at P<5.7×10^−8^ and represents the value below which CpG sites were considered to have good evidence of association with HPV16 E6 seropositivity.

In the DMR analysis of HPV16 E6 seropositivity, 31 unique DMRs pertaining to 158 CpGs and annotating to 38 gene regions were identified (**Figure 3**). The most associated DMR was a region of 13 CpGs found at Chr5:110062343-110062838 (P:4.10×10^−6^), annotating to the *TMEM232* gene region.

### Epigenome-wide association study of OPC survival

#### Model 1

In the single-site analysis of survival (adjusting for age, sex and surrogate variables obtained by SVA (32)), three CpGs mapping to three unique loci showed association with survival below the multiple testing p-value threshold (P<5.7e^−8^) (**Figure 4**). One CpG site showed lower methylation in those who died vs were alive during follow-up. This site was also the most strongly associated with survival, annotating to *PAQR3* and showed the largest effect size among the top hits (cg25864218; β [difference in methylation between those that were dead vs alive before 30^th^ September 2017]: −2.54%; *P:* 1.04e^-9^). Two sites showed higher methylation in those who died vs were alive during follow-up in our analysis, annotating to *DNAH11* (cg07377396; β: 0.49%; *P:* 3.39e^-8^) and *MYBPC1* (cg12151015; β: 0.11%; *P:* 7.51e^-9^). The mean difference in methylation in these sites was 0.3% (SD: 0.27%, range: 0.11% to 0.49%). All results below a suggestive multiple testing threshold of 2.4e^−7^ are shown in **Supplementary Table 3.**

**Figure 4.**
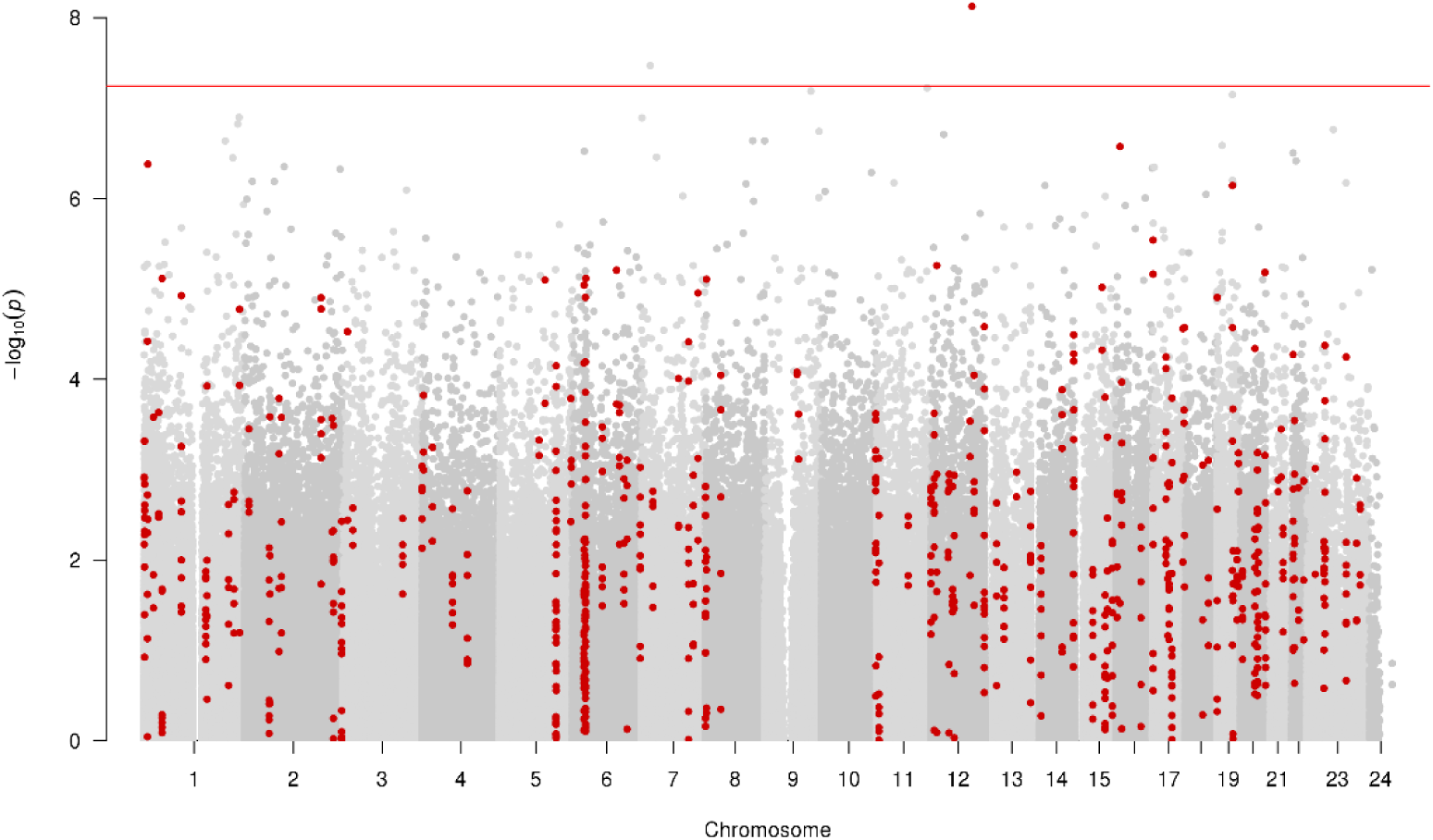
Manhattan plot of EWAS of survival (model 1 – adjusting for age, sex and surrogate variables obtained by SVA), showing CpG sites within DMRs in red. Each dot represents the EWAS result for a single CpG site, plotting –log_10_(p) (y-axis) against the genomic position of the CpG site (x-axis). The horizontal red line is at P<5.7×10^−8^ and represents the value below which CpG sites were considered to have good evidence of association with survival.

In the DMR analysis of survival, 142 unique DMRs pertaining to 805 CpGs and annotating to 153 gene regions were identified (**Figure 5**). The DMR with the lowest P value was a region of 10 CpGs found at Chr17:33814297-33814897 (P:5.26e^-21^), annotating to the *CDK16* gene region.

**Figure 5.**
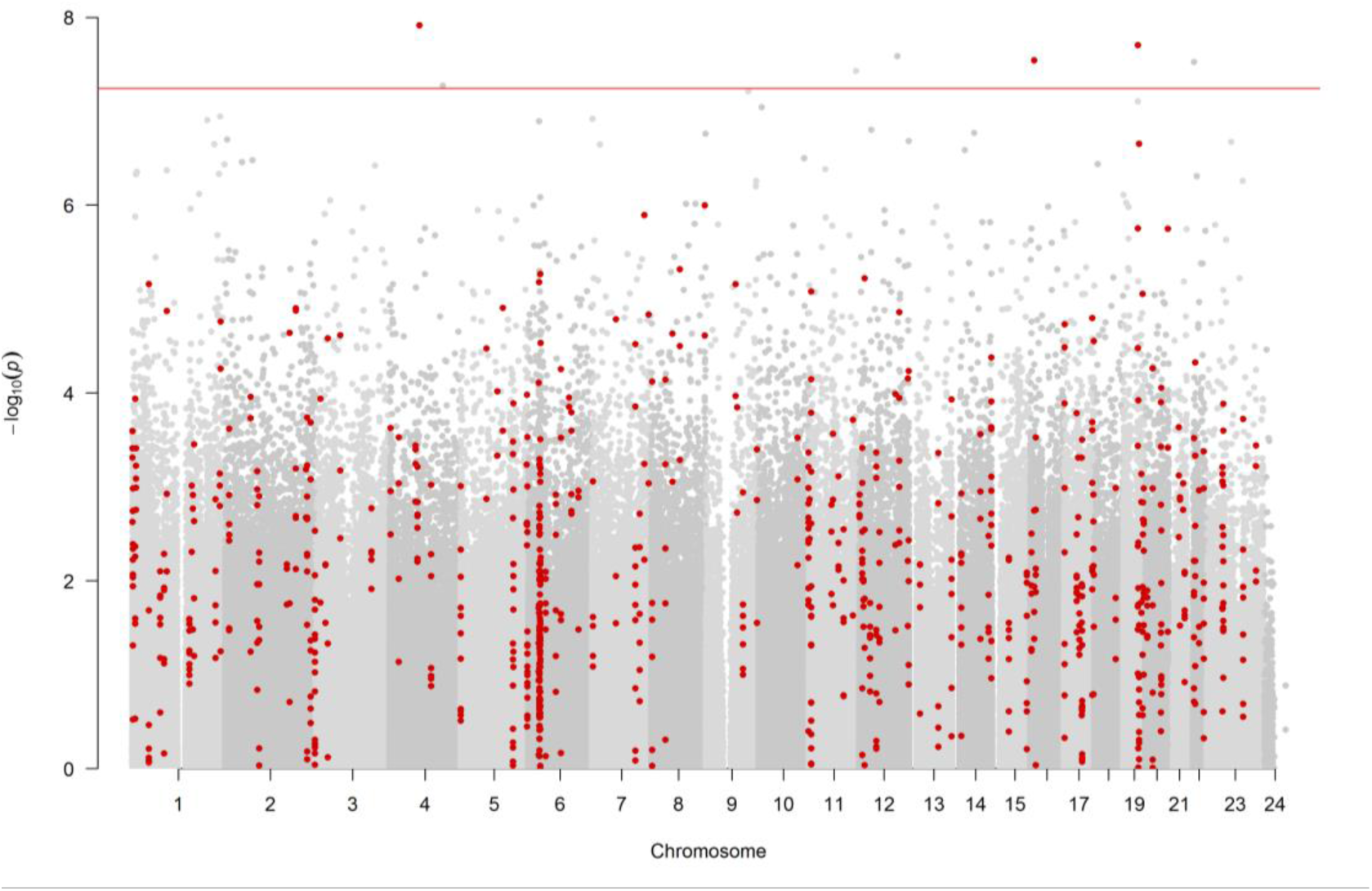
Manhattan plot of EWAS of survival (model 2 – adjusted for smoking, alcohol consumption and HPV16E6 seropositivity), showing CpG sites within DMRs in red. Each dot represents the EWAS result for a single CpG site, plotting – log_10_(p) (y-axis) against the genomic position of the CpG site (x-axis). The horizontal red line is at P<5.7×10^−8^ and represents the value below which CpG sites were considered to have good evidence of association with survival.

#### Model 2

In the single-site analysis of survival using Model 2 (adjusting for age, sex, surrogate variables obtained by SVA (32), HPV16E6 seropositivity, smoking status and alcohol intake), 6 CpGs annotated to 4 unique loci showed a p-value of association below our multiple testing threshold (P<5.7e^−8^) (**Figure 5**). Three of the 6 CpGs passing multiple testing correction showed lower methylation in those who died vs were alive during follow-up in our analysis, while the other 3 showed higher methylation. Of the 3 sites showing lower methylation, the mean difference in methylation between those that were dead vs alive after ∼3-year follow-up was −0.07% (SD: 0.05%, range: −2.54% to −0.16%). For the 3 sites showing higher methylation, the mean difference in methylation was 0.31% (SD: 0.31%, range: 0.11% to 0.67%). The CpG with the smallest P value (cg25864218, *P:* 1.22×10^−8^), annotates to the *PAQR3* gene region. This CpG site also showed the largest effect size of −2.5% difference in methylation between those who are dead vs alive in this analysis. Other CpGs passing our multiple testing correction which were annotated to genes included *MYBPC1* (cg12151015; β: 0.11%; *P:* 2.59e^-8^), *GRIN2A* (cg08204867; β: −0.16%; *P:* 2.87e^-8^), and *IL15* (cg26269613; β: 0.67%; *P:* 5.34e^-8^). Two CpGs showed an association with survival in both models: cg12151015 (annotating to *MYBPC1*) and cg25864218 (annotating to *PAQR3*). All results below a suggestive multiple testing threshold of 2.4e^−7^ are shown in **Supplementary Table 4.** Interestingly, of the results presented in this table, all 23 associated CpGs were present on the EPIC array but not the 450K predecessor.

In the DMR analysis of survival (model 2), 157 unique DMRs pertaining to 874 CpGs and annotating to 177 gene regions were identified (**Figure 5**). The DMR with the lowest P value was a region of 12 CpGs found at ChrX: 47077168-47077877 (P:1.08e^-21^), annotating to the *CDK16* gene region.

### DMR overlap between OPC risk factors and survival

Eighteen unique CpGs overlapped between all smoking DMRs and survival DMRs (survival EWAS model 1). These CpGs belonged to 3 unique DMRs (annotated to *GFI1, SPEG* and *PPT2*); five CpGs overlapped between all alcohol DMRs and survival (EWAS Model 1) DMRs, all pertaining to a single DMR (annotated to *C6orf221*) (**Supplementary Table 5**). No CpGs overlapped at our p-value threshold for HPV DMRs and survival (EWAS model 1) DMRs.

Of the 18 CpGs which overlapped between smoking and survival, 15 possessed mQTL proxies in the Generation Scotland summary data with which to conduct MR (see Methods). Of the 5 CpGs which overlapped between alcohol and survival, 3 possessed mQTL proxies in the Generation Scotland summary data (**Supplementary Table 5**).

### Mendelian randomization: DNA methylation - OPC survival

**Tables 2a-c** and **Figure 8** show the results MR analyses for the association of mQTL-proxied DNA methylation, at CpG sites associated with smoking and survival, with 3-year survival in HN5000. In our analyses, there appears to be some evidence for a potential causal effect of decreased DNA methylation on survival at the *SPEG* gene locus (**Table 2a**; Chr2:22035443-22036041; HR: 1.28; 95% CI: 1.14 to 1.43). Our results provide evidence of a causal association seen between methylation changes in response to smoking, and decreased survival at this gene region. A lookup in the BIOS QTL Browser (https://genenetwork.nl/biosqtlbrowser/) was conducted to assess whether methylation at this locus affected gene expression; twenty cis-expression quantitative trait methylations (eQTMs) showed evidence of correlation between gene expression and methylation at the *SPEG* locus in whole blood at this gene region.

**Table 2a.** Mendelian randomization (MR) analysis results, assessing epigenetic mediation between smoking status and ∼3-year survival at the SPEG gene (chromosome 2:220325443-220326041). The number of SNPs per analysis are shown, in addition to the inverse-variance weighted (IVW) and multivariable MR Egger MR results. IVW and MR Egger results are adjusted for genetic correlation between mQTLs are reported as hazard ratios (HR) with 95% confidence intervals (CI). The SPEG locus was the only in our analyses to possess >2 independent SNPs and is therefore the only with multivariable MR Egger analysis conducted on this independent subset in addition to all DMR CpGs.

| Region (gene) | MR Method | SNPs | HR | 95% CI | P |
| --- | --- | --- | --- | --- | --- |
| **All DMR CpGs** |  |  |  |  |  |
| Chr2:220325443-220326041 (*SPEG*) | IVW | 17 | 1.28 | 1.14 to 1.43 | 2.12x10<sup>-05</sup> |
| Chr2:220325443-220326041 (*SPEG*) | MR Egger | 17 | 1.28 | 1.18 to 1.38 | 4.04x10<sup>-10</sup> |
| **Sentinel CpG only** |  |  |  |  |  |
| cg06084174 (*SPEG*) | IVW | 3 | 1.14 | 0.90 to 1.45 | 0.29 |
| **CpGs with independent SNPs** |  |  |  |  |  |
| Chr2:220325443-220326041 (*SPEG*) | MR Egger | 4 | 1.27 | 0.78 to 2.08 | 0.34 |

**Table 2b.** Mendelian randomization (MR) analysis results, assessing epigenetic mediation between smoking status and ∼3-year survival at the GFI1 gene (chromosome 1:92946132-92947588). The number of SNPs per analysis are shown, in addition to the inverse-variance weighted (IVW) and multivariable MR Egger MR results. IVW and MR Egger results are adjusted for genetic correlation between mQTLs are reported as hazard ratios (HR) with 95% confidence intervals (CI).

| Region (gene) | MR Method | SNPs | HR | 95% CI | P |
| --- | --- | --- | --- | --- | --- |
| **All DMR CpGs** |  |  |  |  |  |
| Chr1:92946132-92947588 (*GFI1*) | IVW | 8 | 0.74 | 0.60 to 0.93 | 7.9x10<sup>-03</sup> |
| Chr1:92946132-92947588 (*GFI1*) | MR Egger | 8 | 2.65 | 0.77 to 9.12 | 0.12 |
| **Sentinel CpG only** |  |  |  |  |  |
| cg06338710 (*GFI1*) | Wald ratio | 1 | 0.93 | 0.47 to 1.85 | 0.84 |

**Table 2c.** Mendelian randomization (MR) analysis results, assessing epigenetic mediation between smoking status and ∼3-year survival at the PPT2 gene (chromosome 6:32120895-32120907). The number of SNPs per analysis are shown, in addition to the inverse-variance weighted (IVW) and multivariable MR Egger MR results. IVW and MR Egger results are adjusted for genetic correlation between mQTLs are reported as hazard ratios (HR) with 95% confidence intervals (CI).

| Region (gene) | MR Method | SNPs | HR | 95% CI | P |
| --- | --- | --- | --- | --- | --- |
| **All DMR CpGs** |  |  |  |  |  |
| Chr6:32120895-32120907 (*PPT2*) | IVW | 8 | 0.82 | 0.52 to 1.30 | 0.40 |
| Chr6:32120895-32120907 (*PPT2*) | MR Egger | 8 | 1.68 | 0.27 to 10.38 | 0.58 |
| **Sentinel CpG only** |  |  |  |  |  |
| cg17113856 (*PPT2*) | IVW | 2 | 0.67 | 0.37 to 1.22 | 0.19 |

**Figure 8.**
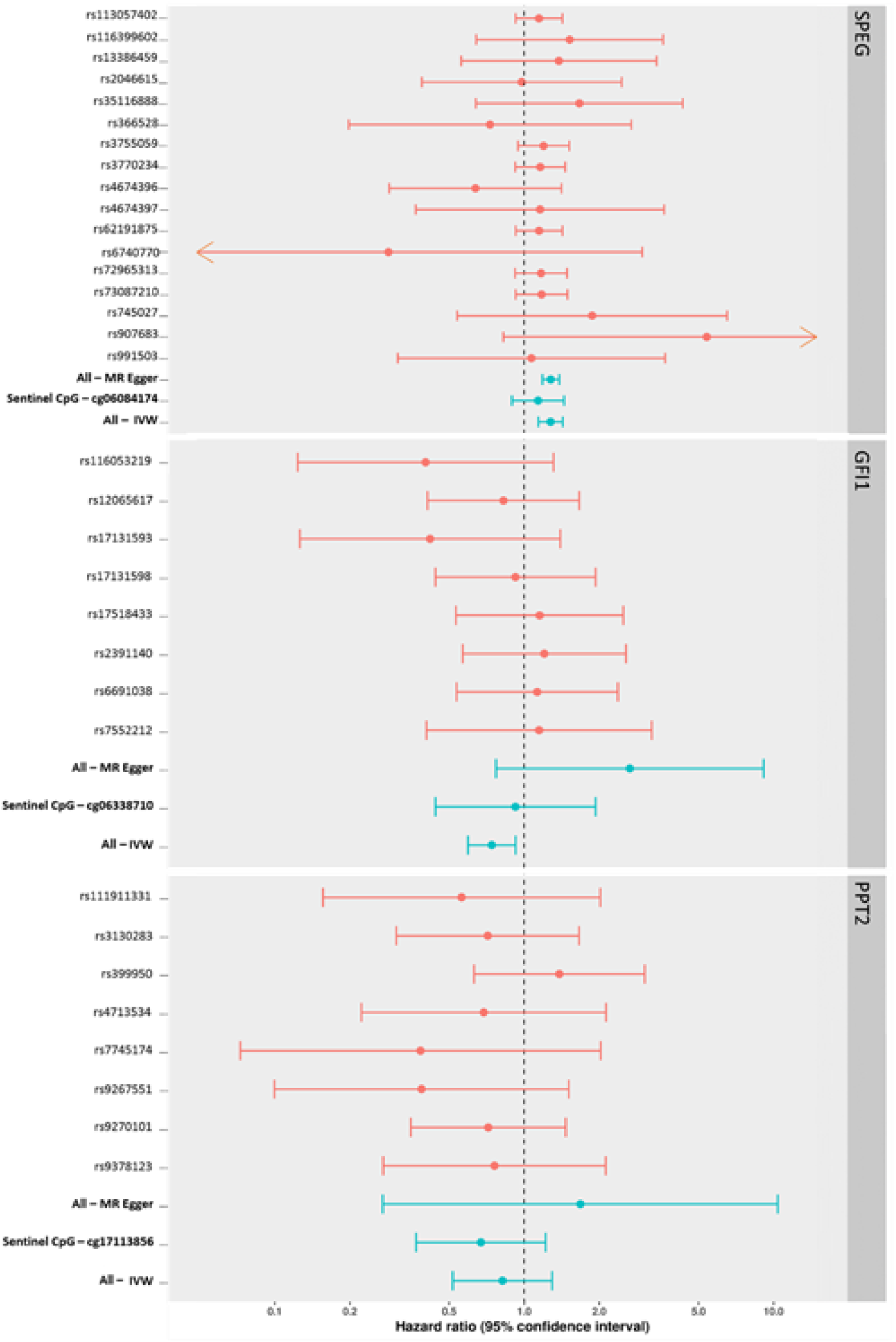
Forest plots showing SNP-specific and overall IV Hazard ratio estimates (95% CI) for Mendelian randomization analyses of smoking-associated methylation at 3 gene loci (GFI1, PPT2, SPEG), against 3-year survival in oropharyngeal cancer.

The *GFI1* (**Table 2b**) and *PPT2* (**Table 2c**) gene regions appear to show no consistent evidence of a causal effect of DNA methylation on survival. We could only conduct multivariable MR Egger analysis using independent SNPs (multivariable MR Egger_independent_: a sensitivity analysis for using multivariable MR Egger with correlated SNPs in our main analysis) at the *SPEG* locus, as other regions did not have sufficient independent SNPs as proxies. Fewer than 3 SNPs greatly reduces the accuracy of MR Egger; therefore, it was only used in analyses with 3 or more SNP proxies. Multivariable MR Egger_independent_ showed a similar effect estimate to normal multivariable MR Egger at this locus, albeit with larger confidence intervals, suggesting the confidence interval for normal multivariable MR Egger is likely to be overly precise in this analysis.

**Table 3** and **Figure 9** show the results of the associations of mQTL-proxied DNA methylation, at CpG sites associated with alcohol and survival with 3-year survival in HN5000. In our analysis, there appears to be no consistent evidence for a causal effect of DNA methylation on survival at the *C6orf221* gene locus (Chr6:74072255-74072376).

**Table 3.** Mendelian randomization (MR) analysis results, assessing epigenetic mediation between alcohol consumption and ∼3-year survival at the KHD3CL gene (chromosome 6:74072255-74072376). The number of SNPs per analysis are shown, in addition to the inverse-variance weighted (IVW) and multivariable MR Egger MR results. IVW and MR Egger results are adjusted for genetic correlation between mQTLs are reported as hazard ratios (HR) with 95% confidence intervals (CI).

| Region (gene) | MR Method | SNPs | HR | 95% CI | P |
| --- | --- | --- | --- | --- | --- |
| **No clumping of final instrument, meta-analysis of mQTLs** |  |  |  |  |  |
| Chr6:74072255-74072376 (*C6orf221*) | IVW | 4 | 1.17 | 0.70 to 1.97 | 0.55 |
| Chr6:74072255-74072376 (*C6orf221*) | MR Egger | 4 | 0.89 | 0.27 to 2.98 | 0.85 |
| **Sentinel CpG only** |  |  |  |  |  |
| cg19146112 (*C6orf221*) | Wald ratio | 1 | 1.17 | 0.54 to 2.53 | 0.68 |

**Figure 9.**
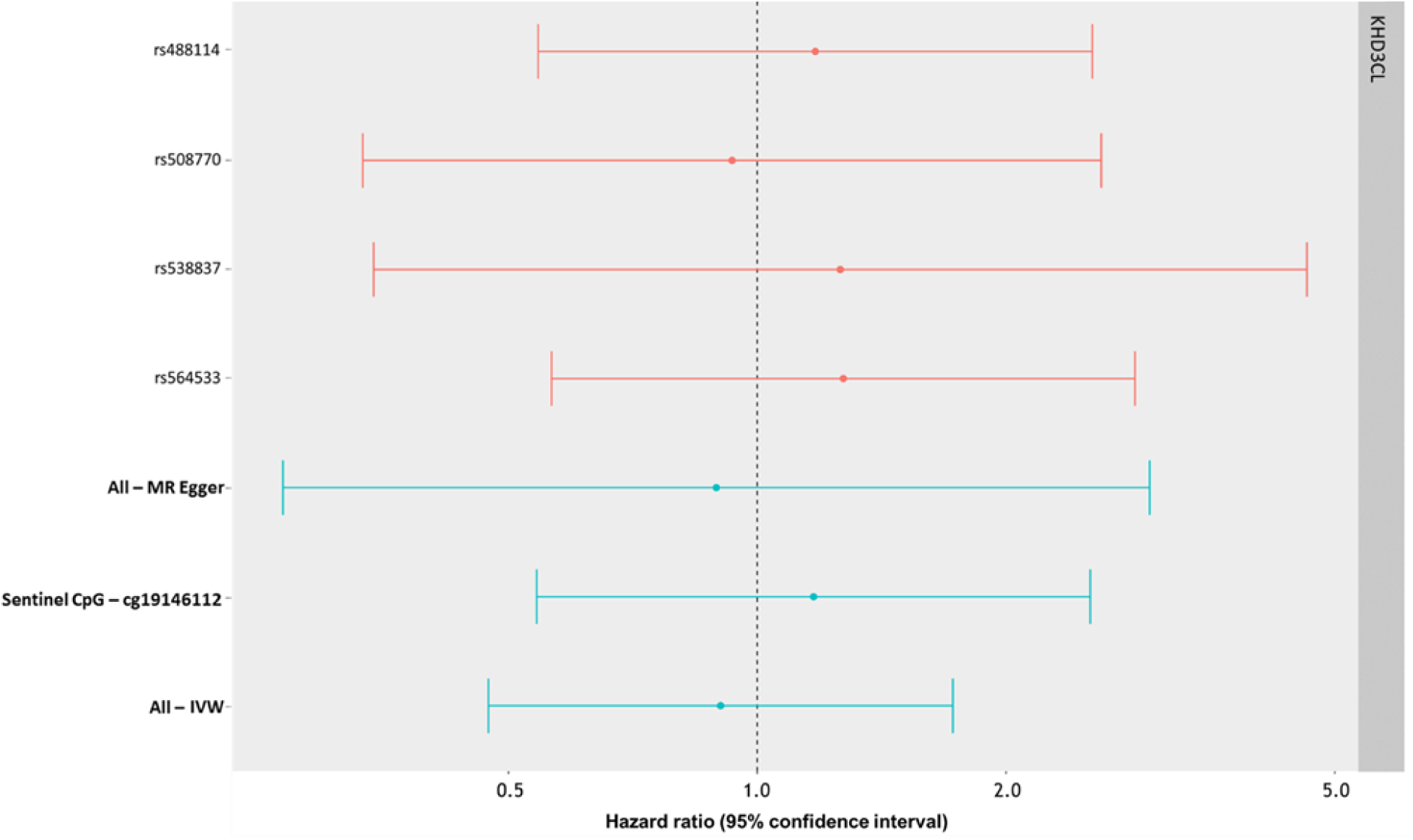
Forest plot showing the SNP-specific and overall IV Hazard ratio estimates (95% CI) for Mendelian randomization analyses of alcohol-associated methylation at the KHDC3L gene locus, against 3-year survival in oropharyngeal cancer.

Sensitivity analyses were conducted for results where MR Egger estimates showed an opposite direction of effect to the IVW estimate (*GFI1, PPT2, KHDC3L*). For each of these analyses, the MR Egger intercept test of heterogeneity (explained elsewhere (33, 34)) spanned our null of 0 (*GFI1* intercept: −0.25, 95% CI: −0.54 to 0.05, p-value: 0.10; *PPT2* intercept: −0.18, 95% CI: −0.58 to 0.23, p-value: 0.40; *KHDC3L* intercept: 0.07, 95% CI: −0.09 to 0.23, p-value: 0.37), providing evidence that directional pleiotropy was not causing the difference between the MR Egger and IVW estimates.

## Discussion

We undertook EWAS analyses and identified CpG sites and DMRs associated with smoking and alcohol consumption, but none associated with HPV infection. We also identified 6 CpGs associated with OPC survival at 3 years post-diagnosis. Twenty-three CpGs at 4 DMRs were identified in both analyses of risk factor and of survival. We hypothesised that for these CpG sites, DNA methylation could mediate part of the association between the risk factor and OPC survival. MR analysis was conducted to test this hypothesis and we found preliminary evidence to support this mediation pathway between smoking and OPC survival at the *SPEG* gene locus.

In relation to smoke exposure, our results include several previously reported loci, notably those mapping to *AHRR* and *PRSS23*. The effect size seen in our EWAS for cg05575921 (29.5%) is markedly stronger than the largest published smoking EWAS analysis; Joehanes et al (21) report 18% lower methylation in current smokers compared to those who have never smoked (P: 4.60e^-26^). A potential explanation for this finding could be that our analysis was conducted in a case-only setting where smoking is one of the predominant risk factors for HNC and smoking intensity is likely to be higher in HN5000 smokers compared to smoking in the general population. We completed a lookup of our top smoking CpG sites (*P* < 5.7e-8), using the EWAS Catalog (http://www.ewascatalog.org/) online tool to compare whether our effect sizes were consistently stronger than other published smoking EWAS findings (**Supplementary Table 6**). Of the 52 sites below a multiple-testing correction, 20 had not been previously reported in published EWASs. The other 32 CpG sites which had previously been reported in the literature showed consistently larger effect estimates in response to smoking in our analysis compared to a weighted mean (weighted by sample size) of published EWAS beta values.

None of the 5 CpG associations with alcohol were replicated in data from the EWAS catalog. However, CpGs associated with alcohol were specific to the EPIC array and no other EPIC array EWASs of alcohol are currently indexed by the EWAS catalog. *SLC7A11*, the gene annotated to our top CpG site associated with alcohol consumption, is essential for glutathione synthesis, a component of the KEAP1-NRF2-CUL3 axis, and strongly associated with poor prognosis in The Cancer Genome Atlas (TCGA) HNC cohort (35, 36).

In our EWAS of 3-year survival none of the 15 (model 1) or 23 (model 2) reported associations have previously been reported in published studies of OPC survival. Both survival EWAS models gave a top hit annotating to the *PAQR3* gene. Aberrant promotor methylation at this gene has been shown to be associated with prostate cancer (37), with the gene itself being an established tumour suppressor (38). Within the context of HNC, *PAQR3* has been associated with tumorigenesis in oesophageal cancer (39, 40), although to our knowledge no literature has examined whether this gene affects oropharyngeal cancer specifically.

The consistent direction of effect between MR Egger, MR Egger_independent_ and Burgess IVW estimates for the *SPEG* locus provide us with greater confidence that the IV is reliable and that there is sufficient statistical power to demonstrate preliminary evidence for a causal association with decreased survival. Expression of the *SPEG* gene shows specificity to vascular smooth muscle cells – the major cell type in blood vessel walls, in which smoking has been shown to produce abnormal function throughout the body (41). Functional annotations show the *SPEG* gene to be essential for cardiac function in particular, with deficiency of this gene reported to result in heart failure (42). As mentioned previously, a lookup in the BIOS QTL Browser (https://genenetwork.nl/biosqtlbrowser/) confirmed 20 eQTMs showing evidence of correlation between gene expression and methylation at this locus in whole blood, though further work evaluating tissue-specific expression is required. People with head and neck squamous cell carcinoma (HNSCC) have an elevated risk of non-HNSCC mortality that persists over their lifetime. Among people with HNSCC, the 5-year incidence of non-cancer mortality is 13% (43), with a high baseline risk of cardiovascular disease compared to matched controls (44, 45).

To our knowledge, this is the first EWAS study investigating oropharyngeal cancer survival using a cox proportional-hazards model to investigate DNA methylation in relation to survival at ∼3 years. This study uses data derived from the EPIC array, which profiles methylation at approximately twice as many CpG sites as its 450k predecessor. Across the EWASs of smoking, alcohol, HPV and both survival models, 39.4% of the CpG sites showing association at *P*<2.4e-7 were specific to the EPIC array (43/109). However, proportionally, our results suggest that associations are not enriched with the inclusion of novel enhancer region CpGs from the EPIC array. A one-sided Fisher’s exact test for enrichment of EPIC probes vs 450K probes in CpG sites below *P*: 2.4e-7 confirms this; *P* > 0.99, suggesting no evidence of enrichment.

The HN5000 study recruited individuals with HNC close to time of diagnosis, taking blood samples prior to treatment, negating potential confounding of methylation changes in response to treatment and minimising information bias. However, whilst unlikely, because cases weren’t recruited prior to HNC diagnosis, we cannot rule out that the differences observed in methylation patterns for smoking (ever vs never), alcohol consumption (units/week) and HPV16 E6 seropositivity (vs HPV16 E6 seronegative) are a result of early stage disease difference. By extension, we cannot state with complete certainty that methylation is an intermediate causal agent; it is possible that a methylation → survival pathway exists independently (i.e. not mechanistically connected) of a smoking → methylation pathway, rather than our hypothesised smoking → methylation → survival pathway.

It should be noted that, despite being an established biomarker with high sensitivity (>93%) and specificity (>94%) for HPV-driven OPC (46), HPV16 E6 seropositivity may underestimate the number of individuals in our data with a current HPV-driven disease; they may be yet to present with the disease. Additionally, it has been reported that HPV can colocalise to biofilm (a community of immotile bacteria encased in a self-produced glycocalyx matrix) in tonsillar crypts, representing a reservoir of latent oncovirus undetected by the immune system (47). Therefore, it is also possible that individuals in our data have a historically HPV16-driven OPC without evidence of infection at time of assessment. As such, our EWAS results for HPV16 infection may be biased towards the null in both instances.

Collider bias may influence associations between our prognostic factors and progression in a case-only setting (48). HPV, smoking and alcohol are all associated with OPC incidence; by only examining cases, incidence is conditioned on, potentially inducing an association between HPV, smoking, alcohol and any unmeasured confounding. Unmeasured, unknown, confounding cannot be adjusted for here, so if any unmeasured confounding is associated with survival, it may be that an association between a prognostic factor and survival is simply a result of the induced association of the prognostic factor and unmeasured confounding.

Some of our MR analyses highlight potential violations of its methodological assumptions. Primarily, those analyses where the MR Egger estimate shows an opposite direction of effect to the IVW estimate (*GFI1, PPT2, KHDC3L*) could indicate an IV where one or more of the genetic variants proxying methylation is disproportionately skewing the effect in a certain direction (horizontal pleiotropy). However, for each of these analyses, the MR Egger intercept test of heterogeneity (explained elsewhere (33, 34)) spans 0 (*GFI1* intercept: −0.25, 95% CI: −0.54 to 0.05, p-value: 0.10; *PPT2* intercept: −0.18, 95% CI: −0.58 to 0.23, p-value: 0.40; *KHDC3L* intercept: 0.07, 95% CI: −0.09 to 0.23, p-value: 0.37), indicating that directional pleiotropy is not causing the difference between the MR Egger and IVW estimates. A possible explanation of this finding, and one that we cannot rule out, is that these analyses suffer from weak instrument bias; a bias where the chance difference in confounders may explain more of the difference in phenotype between genotype subgroups than the instrument, thereby confounding the true causal estimate. Finally, in these three analyses, we cannot state the true direction of effect with confidence, given that our confidence intervals span our null line of Y = 1; this is likely an artefact of low statistical power.

One notable limitation of our MR analysis is that it is likely particularly conservative as we assessed overlap between prognostic factor DMRs and survival DMRs only if they surpassed our multiple correction threshold in both analyses. We adopted this approach (rather than to test corrected prognostic factor DMRs for association with all survival DMRs, only correcting for a number of tests equal to the number of prognostic factor DMRs) to improve confidence that regional methylation was associated with *both* a prognostic factor and survival. In order to reduce the possibility that regional methylation was only associated with a prognostic factor (and only spuriously associated with survival), we may have missed genuine causal mediation at less-stringent p-value thresholds.

## Conclusions

Within the context of OPC, we found novel epigenetic biomarkers measured by the Illumina Infinium EPIC array to be associated with the prognostic factors of smoking and alcohol and with survival. Of these biomarkers, we used overlapping signals between prognostic factor and survival analyses to conduct MR analysis to appraise the causal role of DNA methylation. Using an IVW approach to investigate the causal effect of DNA methylation at the identified sites, we found that a collection of CpGs located within a DMR associated with smoking (located at Chr2:220325443-220326041; annotating to the *SPEG* gene) showed some evidence of a causal effect on decreased survival (HR: 1.28, 95% CI: 1.14 to 1.43, *P:* 2.12×10^−05^). DNA methylation at this locus could potentially mediate some of the association between smoking and OPC survival. To strengthen the validity of these findings, replication analyses in other studies, and a longer follow-up period in Head and Neck 5000 are recommended.

## Methods

### Study population

The study population for this analysis were individuals enrolled in the Head and Neck 5000 (HN5000) clinical cohort study. Full details of the study methods and overall population are described in detail elsewhere (49, 50). Briefly, between April 2011 and December 2014, 5511 individuals with HNC were recruited from 76 centres across the UK. All people with a new diagnosis of HNC were eligible to join the study and were recruited before or within a month of their cancer treatment commencing. Individuals with cancers of the pharynx, mouth, larynx, salivary glands and thyroid were included, while those with lymphoma, tumours of the skin or a recurrence of a previous head and neck cancer were excluded from the study. There were 119 exclusions between recruitment and our data release (v2.3) for the following reasons: withdrawn by study/ineligible (n = 72), patient choice withdrawal (n = 12), and not HNC (n = 35).

Participants for our study were selected from the wider pool of individuals (post-exclusion) in HN5000 (N: 5392) based on an ICD-10 coding (pathological where available, clinical if otherwise) of oropharynx (CO1, CO5, CO9, C10.0-2, C10.3, C10.8 and C10.9; N: 1909/5392), availability of OncoChip genotype data generated previously (N: 1034/1909) (51), baseline questionnaire and data capture information (see below), and the availability of blood samples taken at baseline (prior to treatment; N: 448/1034).

Local research nurses obtained informed consent from individuals, which included agreement to collect, store and use biological samples; obtain samples of stored tissue; carry out genetic analyses and collect clinical information from hospital notes and mortality data through record linkage. Ethics approval for this study was granted by the National Research Ethics Committee (South West Frenchay Ethics Committee, reference 10/H0107/57, 5th November 2010) and approved by the research and development departments from participating NHS Trusts.

### Baseline data collection

Participants completed a series of three self-administered questionnaires at baseline enquiring about: 1) social and economic circumstances, overall health and lifestyle behaviours; 2) physical and psychological health, well-being and quality of life; and 3) past sexual history and behaviours (49). Information on diagnosis, treatment and co-morbidity was recorded on a short data capture form using questions based on a national audit (52). Diagnoses were coded using the International Classification of Diseases (ICD) version 10 (53) and clinical staging of the tumour was derived based on the American Head and Neck Society TNM staging (54).

Research nurses collected a blood sample from all consenting participants (49). These were then sent to the study centre laboratory at ambient temperature for processing. The blood samples were centrifuged at 3500 rpm for 10 minutes and the buffy coat layer used for DNA extraction. Any additional samples from the same participant were frozen and stored at −80°C.

#### Assessment of tobacco, alcohol and HPV infection

Detailed information on tobacco and alcohol history was obtained at baseline via the self-administered questionnaire. Participants were asked about their current smoking and drinking status and their use of tobacco and alcohol products prior to receiving their HNC diagnosis.

Among smokers, information on age at smoking initiation and number of years of smoking was obtained. The questionnaire differentiated between use of cigarettes, hand-rolled cigarettes, cigars and smokeless tobacco, whereby a cigar was considered equivalent to four cigarettes. From this information, participants were dichotomised into ever and never smokers. Ever smokers were defined as those who smoked at the equivalent of at least 1 tobacco product a day per year, or ≥100 cigarettes in their lifetime. Never smokers were those who reported not smoking in any of the questions answered.

Respondents were asked to report their average weekly alcohol consumption of a range of beverage types (wine, spirits, and beer/larger/cider) before they were diagnosed with cancer. From these measures, we derived an average intake of alcohol consumption in units per week.

HPV serologic testing (HPV16 E6, E7, E1, E2, E4, and L1) was conducted at the German Cancer Research Center (DKFZ, Heidelberg, Germany) using glutathione S-transferase multiplex (55). Median fluorescence intensity (MFI) values were dichotomized to indicate HPV16 E6 seropositivity using a cut-off of ≥ 1000 MFI (56). E6 seropositivity is known to be a marker of with a high sensitivity and specificity for HPV16-driven oropharyngeal cancer (57).

#### Study follow-up and survival

Regular updates were received from the NHS Central Register (NHSCR) and the NHS Information Centre (NHSIC) notifying on subsequent cancer registrations and survival among cohort members in the Head and Neck 5000 study. Recruitment for the study finished in December 2014 and follow-up information on survival status was obtained on 30^th^ September 2017, resulting in at least 2.75 years of follow-up for all participants (median: 3.1 years; range: 2.75 to 4.9 years: inter-quartile range: 1.1 years).

### DNA methylation

#### Data generation

Following extraction, DNA was bisulphite-converted using the Zymo EZ DNA Methylation^™^ kit (Zymo, Irvine, CA, USA). Genome-wide methylation data were generated using the Infinium MethylationEPIC BeadChips (EPIC array) (Illumina, USA) according to the manufacturer protocol. The arrays were scanned using an Illumina iScan (version 2.3).

#### Pre-processing

Raw data files (IDAT files) were pre-processed using the R package *meffil* (https://github.com/perishky/meffil/) (58) to perform quality control (QC) and normalisation. Sample mismatches and outliers were identified and removed based on allosome methylation (N: 2 incorrect sex prediction; N: 3 outliers) and 65 genotype probes, which were compared with SNP-chip data from the same individual (N: 3 exclusions). Sample outliers were also identified based on control probe (bisulfite 1 and bisulfite 2) mean outliers (N: 2 exclusions), outliers for the median intensity of the methylated vs unmethylated signal for all control probes (N: 2 exclusions), detection p-value (N: 2 exclusions based on high proportion of undetected probes [>10% of probes failing a detection p-value > 0.01]) and low bead numbers (N: 1 exclusions). Default thresholds for exclusion in *meffil* were used, with 443 samples passing QC. Following QC, Functional normalization was used to separate biological variation from technical variation (59). Data were normalised using 5 control probe principal components derived from the technical probes. The Infinium EPIC array pipeline detects the proportion of molecules methylated at each CpG site on the array. For the samples, the methylation level at each CpG site was calculated as a beta value (*β*), which is the ratio of the methylated probe intensity and the overall intensity and ranges from 0 (no cytosine methylation) to 1 (complete cytosine methylation).

### EWAS

Epigenome wide association study (EWAS) analysis was conducted to identify associations between DNA methylation and 1) alcohol consumption 2) smoking status and 3) HPV16E6 seropositivity. EWAS were conducted in *meffil*, using a linear regression model of DNA methylation regressed on the prognostic factors, adjusting for age, sex, surrogate variables obtained by SVA (32) and the other prognostic factors (e.g. for alcohol intake, adjusting for smoking and HPV16E6).

Of the 443 individuals who passed QC, the number of individuals with complete phenotype data for alcohol intake, smoking status and HPV16E6 seropositivity with which to conduct an EWAS was 409 as of the 2018, version 2.3 release of HN5000 data. All samples possessed information on survival status.

EWASs for survival from recruitment (last participant recruited December 2014) – September 2017 (or time of censoring; whichever occurred first) was conducted using Cox proportional-hazards models using code adapted from the *meffil* R package (58). Two models were assessed: Model 1, adjusting for age, sex and surrogate variables obtained by SVA (32), and Model 2, adjusting for age, sex, surrogate variables obtained by SVA (32), HPV16 E6 seropositivity, smoking status and alcohol intake. Model 1 was run to assess overlap with prognostic factors by not adjusting for them; Model 2, by adjusting for prognostic factors, would provide survival-specific hits independent of them. Death from any cause was used as the failure variable and time to death (or censoring) in days as the time variable. Other prognostic factors for survival include stage and comorbidity. We conducted survival EWAS with these covariates included and found the effect size remained largely unaffected by the addition of stage and comorbidity (Supplementary Table 7). Therefore, we conducted EWAS of survival without stage and comorbidity as covariates.

Due to the large number of tests conducted in our EWAS, we employed a Bonferroni correction to derive a conservative p-value threshold of 5.7 ×10^−8^ (0.05/862491 independent tests) to determine those sites showing strong evidence of association with our risk factor of interest or survival, respectively. We also used the alpha value calculated for the Illumina 450K array (the predecessor to the MethylationEPIC array) as a p-value threshold of 2.4 × 10^−7^ for suggestive evidence of association (31).

### DMR analysis

Adjacent probes on the Illumina arrays are often highly correlated; therefore, differentially methylated regions (DMRs) may reveal regions of DNA where CpGs are associated with risk factors and survival. Following each EWAS we conducted DMR analysis using the *dmrff* R package (60). This analysis identified regions (> 1 CpG site per region) enriched for low P-values (P<0.05), corrected for dependencies between other CpG sites in the DMR and adjusted for multiple testing.

### Generation Scotland methylation quantitative trait loci

DNA methylation can be influenced by genetic sequence variations, such that individual genotypes at a given locus may result in different patterns of DNA methylation due to allele-specific methylation (61-63). Such sites, called methylation quantitative trait loci (mQTLs), can influence the methylation pattern across an extended genomic region (61), and can be used as a proxy for methylation levels in a Mendelian randomization (MR) framework (29).

To generate mQTLs, methylation data from a quality-controlled subset of individuals (N: 5101) from the Generation Scotland: Scottish Family Health Study (64) who had undergone EPIC array DNA methylation profiling, described previously (65), were used. Following measurement of DNA methylation, normalization was performed using the R package *minfi* (66), producing M-values (67) for downstream analysis. Briefly, linear mixed modelling was used to remove potential effects from technical factors, adjusting for both fixed and random effects. Fixed effects included: the top 50 principal components of control probe intensities (explaining 99% of variation in control probe intensities) (68), clinic centre for blood draw appointment, processing batch, year of clinic visit, and Sentrix position (position of the sample on EPIC array slide). Random effects included: blood draw appointment date and Sentrix ID (EPIC array slide). The model converged successfully for 712,595 sites. Outliers from this normalisation with residualized-M-values more than five interquartile ranges from the nearest quartile were removed (69).

A GKFSC model (70, 71) was then fitted to derive mQTLs from the normalised data, including 5 matrices as random effects, and other covariates as fixed-effects. The matrices were: G (a genomic relationship matrix), K (a kinship relationship matrix) (72, 73), F (an environmental matrix representing nuclear-family-member relationships), S (an environmental matrix representing full-sibling relationships) and C (an environmental matrix representing couple relationships) (70, 71). Covariates (as fixed effects) included: age, age^2^, gender, estimated cell counts, season of clinic visit, appointment time of the day and appointment day of the week. The model successfully converged for 638,737 CpG sites.

### Generation of instrumental variables for DMRs

Prior to MR analysis being conducted (see below), we generated instrumental variables (IVs) proxying CpG sites identified in analyses of both prognostic factors and survival (**Supplementary Figure 1**). Where possible, we found DMRs (*P* < 0.05) from our analyses for each prognostic factor and located DMRs spanning the same region in our survival analysis (Model 1 – unadjusted for prognostic factors; *P* < 0.05). CpG sites present in both DMRs were retained.

Next, using the summary genetic data for mQTLs from Generation Scotland, we extracted all mQTLs proxying any CpG site per DMR grouping (MAF >0.05; *P* < 5×10-8). From this list, we generated instruments by LD pruning iteratively; first taking all mQTLs associated with the sentinel CpG (defined as the CpG in each DMR with the lowest p-value) and clumping with an r^2^ of 0.01. We then took the second most-associated CpG in the DMR and extracted all mQTLs associated with it which were not associated with the previous CpG. The remaining mQTLs were then clumped and combined with the mQTLs proxying the sentinel CpG. This process was repeated for each CpG within a DMR. Clumping and mQTL extraction were conducted using R 3.4.1, with the *TwoSampleMR* R package (74).

In order to account for mQTL proxies influencing methylation at multiple CpG sites, we conducted a meta-analysis of mQTL-CpG effects. Per DMR, we used the *metafor* R package (75) to meta-analyse each mQTL effect (beta) on methylation levels at each CpG using a restricted maximum likelihood (REML) model, adjusting for pairwise correlation between the CpG sites proxied by our instrument. From this, we obtained an mQTL effect on average methylation levels across the DMR.

### mQTL associations with survival

The mQTLs identified above were then regressed against survival in HN5000, using the SurvivalGWAS_SV program in Linux to run Cox proportional-hazards survival analyses with an additive dosage model for each of the selected SNPs (76). Death from any cause was used as the failure variable and time to death (or censoring) in days as the time variable. Age at cancer diagnosis and sex were used as covariables in the model. For each SNP the log-hazard ratio (and standard error) per minor allele was reported.

### Mendelian randomization analyses

Following identification of shared methylation patterns between prognostic factors and OPC survival, we attempted to ascertain whether methylation was a true causal intermediate, or simply just associated with both prognostic factors and survival. To this end, we conducted two-sample Mendelian randomization to appraise the causal effect of DNA methylation on survival. In the first sample, we used mQTL-DMR effect estimates (βGP) from Generation Scotland and in the second sample, mQTL-survival estimates (βGD) from HN5000. For each mQTL, we calculated the log HR per unit (β) increase in DNA methylation at the DMR by the formula βGD/βGP (Wald ratio). Standard errors were approximated by the delta method. Where multiple mQTLs were available for one DMR, these were combined in a fixed effects meta-analysis after weighting each ratio estimate by the inverse variance of their associations with the outcome (IVW approach). In order to account for correlation between mQTLs, we adjusted for genetic correlation using LDMatrix (77) to generate a genetic correlation matrix (1000 Genomes reference standard (78)) of mQTLs, which was included as a covariate in our MR regression analysis (79). In addition to our main analysis detailed above, we conducted multivariable MR Egger analysis as an assessment of IV heterogeneity using the MendelianRandomization R package (80). We also conducted sensitivity MR analyses by calculating the log HR per unit increase in DNA methylation for the sentinel CpG within each DMR we analysed. As above, Wald ratios were calculated for CpGs proxied by a single mQTL and IVW MR estimates were calculated when multiple mQTLs were available to proxy a CpG. Finally, where possible, we conducted multivariable MR Egger analysis on a subset of independent SNPs for each DMR as a sensitivity analysis for using multivariable MR Egger with correlated SNPs in our main analysis.

## Acknowledgements

This publication presents data from the Head and Neck 5000 study. The study was a component of independent research funded by the National Institute for Health Research (NIHR) under its Programme Grants for Applied Research scheme (RP-PG-0707-10034). The views expressed in this publication are those of the author(s) and not necessarily those of the NHS, the NIHR or the Department of Health. Human papillomavirus (HPV) serology was supported by a Cancer Research UK Programme Grant, the Integrative Cancer Epidemiology Programme (grant number: C18281/A19169).

RL, RR, NK HRE, TD, TG, GDS, MS and CR work in the Medical Research Council Integrative Epidemiology Unit at the University of Bristol which is supported by the Medical Research Council and the University of Bristol (MC_UU_00011/5).

RL, RR, NK and CR are supported by a Cancer Research UK program grant (C18281/A19169).

RL is supported by a Cancer Research UK Research PhD studentship (C18281/A20988).

TD is supported by a Wellcome Trust PhD grant (201268/Z/16/Z).

RMW acknowledges salary support from a Wellcome Trust Strategic Award “STratifying Resilience and Depression Longitudinally” (STRADL) (Reference: 104036/Z/14/Z), which funded the profiling of DNA methylation in Generation Scotland: Scottish Family Health Study participants. Generation Scotland received core support from the Chief Scientist Office of the Scottish Government Health Directorates (Reference: CZD/16/6) and the Scottish Funding Council (Reference: HR03006). RMW is an associate member of The University of Edinburgh Centre for Cognitive Ageing and Cognitive Epidemiology (CCACE), part of the cross-council Lifelong Health and Wellbeing Initiative (MR/K026992/1). Funding for CCACE from the Biotechnology and Biological Sciences Research Council and Medical Research Council is gratefully acknowledged.

Generation Scotland received core support from the Chief Scientist Office of the Scottish Government Health Directorates (CZD/16/6) and the Scottish Funding Council (HR03006). Genotyping and DNA methylation profiling of the Generation Scotland samples was carried out by the Genetics Core Laboratory at the Wellcome Trust Clinical Research Facility, Edinburgh, Scotland and was funded by the Medical Research Council UK and the Wellcome Trust (Wellcome Trust Strategic Award “STratifying Resilience and Depression Longitudinally” ((STRADL) Reference 104036/Z/14/Z)). CH was supported by the Medical Research Council (Reference MC_UU_00007/10). AB was supported by a Wellcome Trust funded ECAT fellowship (reference 204979/Z/16/Z). The authors are extremely grateful for the provision of IlluminaMethylationEPIC mQTL data from this study, particularly to: Andrew Bretherick, Yanni Zeng, Rosie M Walker, Toni-Kim Clarke, Chris Haley, Andrew M McIntosh, Kathryn L Evans and Alison Murray.

